# Taming the Late Quaternary phylogeography of the Eurasiatic wild ass through ancient and modern DNA

**DOI:** 10.1101/090928

**Authors:** E. Andrew Bennett, Sophie Champlot, Joris Peters, Benjamin S. Arbuckle, Silvia Guimaraes, Mélanie Pruvost, Shirli Bar-David, Simon S.M. Davis, Mathieu Gautier, Petra Kaczensky, Ralph Kuehn, Marjan Mashkour, Arturo Morales-Muñiz, Erich Pucher, Jean-François Tournepiche, Hans-Peter Uerpmann, Adrian Bălăşescu, Mietje Germonpré, Can Y. Gündem, Mahmoud-Reza Hemami, Pierre-Elie Moullé, Aliye Ötzan, Margarete Uerpmann, Chris Walzer, Thierry Grange, Eva-Maria Geigl

## Abstract

Taxonomic over-splitting of extinct or endangered taxa, due to an incomplete knowledge of both skeletal morphological variability and the geographical ranges of past populations, continues to confuse the link between isolated extant populations and their ancestors. This is particularly problematic with the genus *Equus.* To more reliably determine the evolution and phylogeographic history of the endangered Asiatic wild ass, we studied the genetic diversity and inter-relationships of both extinct and extant populations over the last 100,000 years, including samples throughout its previous range from Western Europe to Southwest and East Asia. Using 229 bp of the mitochondrial hypervariable region, an approach which allowed the inclusion of information from extremely poorly preserved ancient samples, we classify all non-African wild asses into nine clades that show a clear phylogeographic structure revealing their phylogenetic history. This study places the extinct European wild ass, *E. hydruntinus*, the phylogeny of which has been debated since the end of the 19^th^ century, into its phylogenetic context within the Asiatic wild asses and reveals recent gene flow events between populations currently regarded as separate species. The phylogeographic organization of clades resulting from these efforts can be used not only to improve future taxonomic determination of a poorly characterized group of equids, but also to identify historic ranges, interbreeding events between various populations, and the impact of ancient climatic changes. In addition, appropriately placing extant relict populations into a broader phylogeographic and genetic context can better inform ongoing conservation strategies for this highly endangered species.

## Introduction

Human-driven climate change and habitat reduction of a large number of species has led to what is proposed to be the 6^th^ mass extinction of species and a new era termed the “Anthropocene”. This concerns many plant and animal species including a number of emblematic large vertebrates. To appreciate the extent of these recent changes it is useful to develop a long-term perspective of how previous climate oscillations and human impact have influenced species composition and distribution in the past (e.g., [1]). To acquire a deeper insight into population dynamics in the light of climate change and human activities, we used ancient and modern DNA to study the Asiatic wild ass *(Equus hemionus)*, once widely distributed over a large geographical area including most of Asia and Europe. Although closely related to the domesticated donkey (*Equus asinus*), the Asiatic wild ass has never been domesticated. This taxon thus followed a different evolutionary path, witnessing the extinction of several subspecies since prehistoric times while leaving nearly all those remaining endangered. Genetic and phenotypic analyses of present-day individuals tend to overestimate the differences between relict populations, leading to taxonomic over-splitting. Ancient DNA (aDNA) data has the potential to place the genetic diversity of these small disconnected populations into a broader context of diversity and gene flow prior to the modern situation of diminished and geographically isolated populations. This helps to put into perspective the taxonomic divisions necessary for effective decision-making in conservation management.

Although palaeontology has accumulated much data concerning equid evolution, long considered a paradigmatic evolutionary model [2], much taxonomic uncertainty remains. Indeed, the classification of ancient equids based on osteomorphometry is ambiguous since modern skeletons used for comparisons represent mosaics of various, restricted combinations of a relatively small number of characteristics [3]. Whole skeletons are required for reliable morphological determination and these are extremely rare in the fossil record [4]. Consequently, knowledge of past morphological diversity within and between equid species is scarce. These features put the presently accepted equid taxonomy on shaky ground and question interpretations about the ancient geographical and temporal distribution of this taxon ([5], and citations therein).

Unfortunately, distinct populations of Asiatic wild asses are already on their way to extinction before having been well studied [6]. In many high-altitude plains or deserts of Asia, these arid-adapted and cold-tolerant animals have long been the largest and most widespread herbivore taxon, and its disappearance eliminates a major ecological factor in these extreme environments. In contrast to the caballoids, or horses, Asiatic wild asses belong to the stenonids, a group which also includes zebras and the African wild ass along with its domestic form. Currently, Asiatic wild asses are subdivided into two species, *Equus kiang* and *Equus hemionus* with four living and one extinct subspecies, i.e., *E. h. hemionus* (also known as *E. h. luteus*) –the Mongolian kulan or dziggetai; *E. h. khur* - the Indian khur, *E. h. kulan* - the Turkmen kulan, *E. h. onager* –the Iranian or Persian onager, and *E. h. hemippus* –the extinct Syrian wild ass [3,6,7] (see Fig. S9, supporting information, for their geographic ranges). The vast former range of the Asiatic wild ass has undergone a dramatic reduction so that only a few and fragmented populations remain. The two largest surviving populations, the dziggetais in the Mongolian Gobi Desert [6] and the kiangs of the Tibetan plateau [8], still occur over large parts of their former distribution range. However, increased livestock grazing, fencing, construction of railways and highways and poaching also threaten the future of dziggetais and kiangs. The Iranian onagers, the Turkmen kulans, and the Indian khurs are reduced to small pocket populations with contracted distributions in protected areas located either in the endemic centers or in refuge zones in Iran, Turkmenistan and North West India, respectively [6,9,10].

The European wild ass, termed *E. hydruntinus*, or hydruntine, is known only from skeletal remains and prehistoric engravings such as that found in the cave of *Les Trois Frères* in France (Fig 5). The oldest Western European remains that have been attributed to this morphotype are from France and date to around 350,000 years ago (ya) [11]. The hydruntine was widespread during the Late Pleistocene, with a geographic distribution from Western Europe to the Volga, Turkey, the Levant and the Northern Middle East [12-14]. During the Holocene hydruntine populations declined and were reduced to small patches of their previous range, before eventually becoming extinct [13].

Understanding the evolution as well as past and present genetic diversity of these species is essential for the design of appropriate conservation strategies [15]. Asiatic wild asses, however, are not well characterized genetically. The profound lack of data on the past and recent distribution and population structures of these regionally endangered animals is particularly worrisome at a time so critical for the conservation of Asiatic wild asses, and should therefore be addressed quickly in order to define and implement adequate conservation biology strategies.

Paleogenetic analyses of the mitochondrial and, very recently, nuclear genomes preserved in equid bones have allowed researchers to revisit equid taxonomy, which has reduced the number of species proposed in paleontological studies [16-20]. These recent paleogenetic studies suggested that the “oversplitting” of earlier palaeontological work was the consequence of an underestimation of the morphological plasticity of equids throughout their ranges and evolutionary history [19]. Indeed, ancient DNA research has the potential to unravel the phylogeographic structure of populations and species, past migrations, gene flow, erosion of past diversity and population fragmentation. By correctly identifying the past geographic distribution of genotypes, it is possible to reconstruct the sequences of such events (e.g., [21,22].

In order to characterize the ancient and extant genetic diversity and population structure, we studied the mitochondrial lineages of the wild asses from Europe and Asia, over the last 100,000 years from 70 sites in Europe and Asia. We targeted a 295 bp region in the *E. hemionus* mitogenome that encompasses a specific 28-bp-deletion, absent in other equids, which is a useful bar code for this taxonomic group of *Equus* [19]. This approach, making use of the high copy, neutral, matrilineal marker allowed us to include a large number of important ancient samples from warm environments, many having extremely poor DNA preservation. We analyzed 189 archaeological bone and teeth specimens that had been assigned to *E. hemionus sp.* or *E. kiang* dated between 3,500 and 100,000 ya. These samples originated from 49 archaeological sites in ten European and six Southwest Asian countries (Fig. S9 and Table S1). 64 of the archeological samples yielded DNA sequences. In addition, we analyzed 11 historical museum specimens (between 60 and 180 years old) of onagers, hemippi, khurs and kiangs and 53 present-day samples, 94% of which originated from wild individuals, coming from the Gobi Desert and protected nature reserves in Iran and Israel. We show that during the Upper Pleistocene the distribution of the Asiatic wild ass ranged from Western Europe, where it is now extinct, to East Asia where it is still found at present. The genetic relationship between these taxa (see below) explains why we subsume these populations under the term “Eurasiatic wild ass”. We explored the patterns of the past and present genetic diversity to reconstruct the population structure of the species and its evolution since the Late Pleistocene.

## Materials & Methods

Samples used in this study and their archeological contexts are described in the Supporting Information and listed in Table S1.

### DNA extraction

Modern and historical specimens were processed in a laboratory of the Jacques Monod Institute (IJM) dedicated to modern, non-amplified DNA analysis. Ancient specimens (those older than 150 years) were processed in the core facility of palaeogenomics of the IJM, a high containment laboratory physically separated from the modern DNA laboratories and dedicated to the analysis of ancient DNA.

#### Samples from living specimens

Genomic DNA isolation from blood, muscle and lung tissue was performed with the NucleoSpin^®^-Tissue Kit (Macherey-Nagel, Dueren, Germany). Feces were processed using the QIAamp DNA Stool Mini Kit (Qiagen, Hilden, Germany). Skin, hoof, and hair samples were digested with lysis buffers containing 100 mM Tris-HCL pH 8, 3 mM CaCl2, 2% N-lauroyl-sarcosyl, 40 mM DTT, 250 ug/mL proteinase K, 100 mM NaCl2 (modified after [23]) and DNA was subsequently extracted with the NucleoSpin-Tissue Kit (Macherey-Nagel).

DNA extraction from the blood samples from the individuals from the Hai-Bar Yotveta breeding core, Israel, was performed at the Jacob Blaustein Institutes for Desert Research, Ben-Gurion University of the Negev, Israel, using QIAamp DNA Mini Kit (QIAGEN, Cat. No. 51304) following the manufacturer’s instructions.

#### Samples from museum specimens

Museum samples (the oldest being 174 years old) were processed in the laboratory facilities dedicated to analysis of modern, non-amplified DNA, which is physically separated from the ancient DNA facility and post-amplification laboratory, using aDNA procedures. Sinew and cartilage samples were crushed in liquid nitrogen in a mortar and DNA was purified using the QIAamp DNA Stool Minikit (Qiagen, Hilden, Germany).

#### Ancient Samples

Ancient samples were processed in the Core Facility of Palaeogenetics at the IJM, Paris (http://www.ijm.fr/ijm/plates-formes/pole-paleogenomique/). This highly contained pressurized laboratory dedicated to aDNA analysis is isolated on the 6^th^ floor of the institute where no other laboratories of molecular biology are located. It consists of an airlock chamber and three different laboratory rooms, each subjected to a positive air pressure gradient, dedicated to the specific steps of the experimental procedures, (i) sample preparation, (ii) DNA extraction and purification, (iii) PCR setup. Within the laboratory, each experimental step is carried out in a working station or flow hood. The working stations and equipment are cleaned with bleach (3.5% hypochlorite solution) or RNase away^®^ (Molecular Bio-Products, USA) and UV irradiated at short distance between each experiment to ensure efficient decontamination [24]. To minimize contamination with exogenous DNA and maximize the aDNA yield, low retention microtubes (Axygen, Union City, USA) and extra-long filtered pipette tips were used for extraction and PCR preparation. Experimenters entered the laboratory only after having removed their street clothes and replaced them with lab clothing washed with bleach. During each step of pre-PCR work an all-body protection was worn (a different one for each laboratory room) consisting of a disposable protective suit, two pairs of gloves, shoe covers and a facemask. Purification was carried out in a flow hood and pipetting of the PCRs in a template tamer that were both bleached and UV-irradiated at a short distance after each experiment. Capillaries containing the reagent mixtures for the PCR and the fossil extracts were closed in the template tamer of the PCR-preparation room of the contained laboratory prior to transfer to the PCR machines in a laboratory on the 5^th^ floor of the building. The handling of post-PCR products was exclusively performed on the 5^th^ floor of the building in a dedicated laboratory that is physically separated from the pre-PCR laboratory and the room containing the PCR machines. Protective disposable clothing and shoe covers are also worn when entering the post-PCR laboratory and removed when exiting.

Bone and tooth surfaces were removed with a sterilized razor blade and, depending on the individual bone characteristics, either drilled with a heat-sterilized bit using either a Dremel 4000 or Dremel Fortiflex (Dremel Europe, The Netherlands), or cut into fragments and powdered using a freezer mill (SPEX CertiPrep 6750, USA), or powdered by hand using a razor blade. Bone powder (34-1896 mg) was incubated in 1-10mL extraction buffer (0.5 M EDTA, 0.25 M PO_4_^3-^, pH 8.0, 1% beta-mercaptoethanol) for 48-70 hours at 37°C with agitation. Some samples required changes of extraction buffer to increase digestion of bone powder. Hair and hoof samples were added to a hair digestion buffer (100 mM Tris-HCl pH 8.0, 100 mM NaCl, 40 mM DTT, 3 mM CaCl2, 2% N-lauryl sarcosyl, 250 μg/mL proteinase K) and incubated 4-24 hours at 50°C, shaken with 300 RPM. Solutions were then pelleted and the supernatant was purified using a QIAquick Gel extraction kit (Qiagen, Hilden, Germany) with protocol modifications [25].

For external reproduction of the results, some ancient samples (S1, D4, K2, ASE24, ASE25, ASE382, ASE383, ASE384) were analyzed by M.P. in the laboratory of the Humboldt-University Berlin, Landwirtschaftlich-Gartnerische Fakultat, Molekularbiologisches Zentrum Ostbau, Berlin, Germany

Approximately 250 mg of bone material was used for each extraction. External surfaces of bones were removed by abrasion to minimize environmental contaminations. Each sample was ground to powder with a freezer mill and incubated in 0.45 M EDTA (pH 8.0) and 0.25 mg/ml Proteinase K overnight at room temperature under rotation. After centrifugation for 5 min at 4,000 rpm in a Universal 320 centrifuge (Hettich), DNA was purified from the supernatant using a silica based method as previously described [26,27].

### DNA amplification and sequencing

Primers for archeological samples were designed to amplify up to 295 base pairs of the hyper variable region of wild ass mitogenome using short, overlapping fragments (see Table S2 for a list of primers used and product sizes).

Purified DNA was amplified by qPCR, the extract making up 5-20% total reaction volume (10-20μl). Inhibition characteristics were determined for failed samples indicating possible inhibition. For evaluation of the inhibition, three parameters were taken into consideration: (i) delay of the threshold value Ct (crossing point at threshold), (ii) the kinetics of the synthesis of the PCR product, and (iii) the efficiency of the PCR. The quantity of aDNA extract amplified by PCR was adjusted according to the results of the inhibition tests to minimize interference of the inhibitors with the PCR.

To protect against cross-contamination, the UQPCR method was used [24,28] in which uracil was substituted for thymidine for all PCRs, and incubation with uracil N-glycosylase (UNG, extracted from *G. morhua*; ArcticZymes, Tromso, Norway) was performed prior to each reaction. Mock extraction blanks were performed with each extraction and amplified to control for contamination. qPCR reactions varied slightly depending on the sample, but a typical reaction included 1.77 μL of LC FastStart DNA MasterPLUS mix1b and 0.23 μL of either FastStart DNA MasterPLUS mix 1a or FastStart Taq (Roche Applied Science, Mannheim, Germany), 1 μM of each primer and 1U of UNG and water to 10 μL total volume. Primers were obtained from Sigma-Aldrich (St. Louis, USA), and were designed to amplify 357 base pairs of the hyper variable region of hemione mitochondria using short, overlapping fragments. A list of primers used and product sizes is given in Table S2.

For each ancient sample two independent extractions were performed, separated in time. Each extract was amplified with several primers and each PCR product was obtained at least twice. An average of 1 non-template control (NTC) was run for every 6.6 samples (including mock extracts). No DNA was amplified in either NTCs or mock extracts, indicating no detectable equid DNA was introduced during sample or PCR preparation, or was present in reagents.

qPCR was performed using Lightcycler 1.5 or Lightcycler 2 (Roche Applied Science, Mannheim, Germany). QPCR programs varied depending on primer requirements and product length, but a typical program involved UNG incubation at 37°C for 15 minutes, followed by polymerase activation at 95°C for 8 minutes, 60-80 2-step cycles of denaturation at 95°C for 10 seconds, primer annealing and extension at 62°C for 40 seconds, and finally a temperature increase of 0.1°C/1second from 62°C to 95°C with continuous fluorescence measurement to generate melt-curves of the products. Products were purified by a QIAquick PCR purification kit (Qiagen, Hilden, Germany) and both strands were sequenced by capillary electrophoresis at Eurofins/MWG Operon (Ebersberg, Germany) using an ABI 3730xl DNA Analyzer (Life Technologies). Sanger electropherograms were visually inspected and sequences manually curated, assembled and aligned using the Geneious software suite [29]

PCR analyses of the samples replicated at Humboldt University in Berlin were performed using an overlapping set of primers as described [30]. Using these primers, 713 bp (15.468-16.181 nps) were amplified with a two-step multiplex PCR. Overlapping PCR products, including primers, varied in length between 108 bp and 178 bp. PCR conditions were as described [30]. Multiple negative extraction controls and negative PCR controls were performed. Amplification products were visualized on agarose gel and sequenced on an ABI PRISM 3730 capillary sequencer using the BigDye Terminator v3.1 cycle sequencing kit (Applied Biosystems). The resulting sequences were aligned with those obtained in Paris and found to be identical over their overlapping lengths.

DNA amplification from blood samples from the individuals from the Hai-Bar Yotveta breeding core, Israel, was performed at the Jacob Blaustein Institutes for Desert Research, Ben-Gurion University of the Negev, Israel. The 20 μl reactions contained 20 ng DNA, 0.25 μM of each of the primers EA1 and EA63, 500 μM dNTPs, 2 mM MgCl2 and 1 U Taq DNA polymerase (Hylabs, Rehovot, Israel). PCR conditions were: 5 min at 95°C followed by 40 cycles of 10 sec at 95°C, 30 sec at 62°C and 15 sec at 68°C, and a final elongation step of 10 min at 72°C. PCR products were purified using Exosap IT (USB, Cleveland, OH), and sequenced by ABI 3730XL DNA analyzer (Applied Biosystems).

### Phylogenetic and phylogeographic analyses

Since several of the archeological samples did not yield the complete 295 bp region (see Fig. S1), we selected a 229 bp sequence for phylogenetic analyses that was obtained from 26 archeological samples, nine museum specimens, 52 modern samples, supplemented with the 17 sequences currently found in public databases. In total, we generated 114 new HVR sequences of the Eurasiatic wild asses.

#### Spatial genetics (sPCA)

To investigate the potential relationship between geographic distances and genetic variation, a spatial principal component analysis (sPCA) was carried out on all available sequence information of georeferenced samples using the R packages ade4 and adegenet [31-33].

While with PCA the optimization criterion only deals with genetic variance, sPCA aims at finding independent synthetic variables that maximize the product of the genetic variance and spatial autocorrelation measured by Moran’s *I* [34]. This is accomplished by the eigenvalue decomposition among individuals via a neighboring graph (in this study a Delaunay triangulation was chosen) connecting the individuals on the geographical map to model spatial structure. Resulting eigenvalues can be either positive or negative reflecting respectively either a global or local spatial pattern. A global structure implies that each sampling location is genetically closer to neighbors than randomly chosen locations. Conversely, a stronger genetic differentiation among neighbors than among random pairs of entities characterizes a local structure. To evaluate the consistency of the detected geographical structures versus a random spatial distribution of the observed genetic variance, a Monte-Carlo based test was applied [33]. This test simulated a random distribution of the genetic variability (null hypothesis) on the Delaunay triangulation connection network and calculated a *p*-value depending on the dataset. The simulated distribution represents the correlation of the randomized genetic variables with the vectors of the Moran’s *I* predicting for the global or local structure. If the value associated to the observed pattern is higher than the *p*-value, it means that the spatial distribution of the genetic variance is not random and the null hypothesis can be rejected. We applied the test with 10,000 iterations.

For representing the sPCA scores of each individual we used the colorplot function implemented in the R-software package adegenet [32]. This function can show up to three scores at the same time by translating each score into a color channel (red, green, and blue). The values obtained were expressed as a color under the RGB system as a parameter of the individual genetic constitution and were projected on a world map.

For the representation of the individual sequences on the geographical map in Fig. 1 and to show all samples analyzed, each individual sample was placed on the map as closely as possible to its geographical location and colored using the attributed RGB values of the sPCA scores. The colors obtained with the sPCA analysis were also manually assigned to each sequence on the Beast tree in Fig. 3.

**Figure 1:**
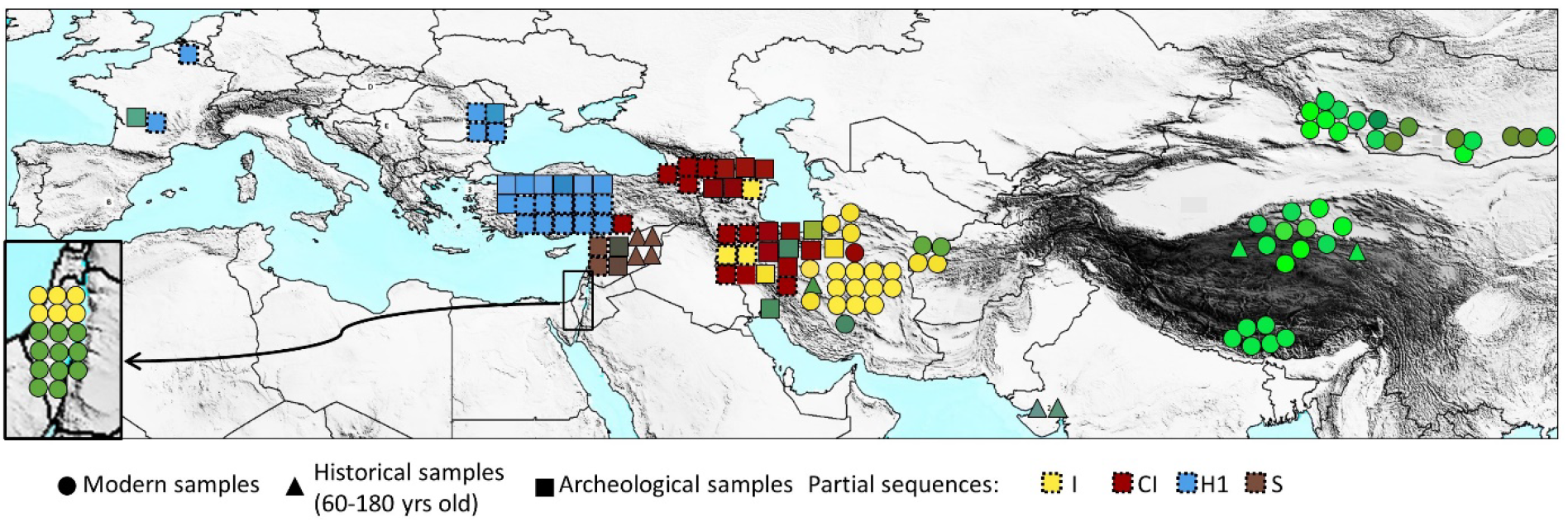
Map representing the results of the landscape genetics sPCA analyses of the samples. Spatial PCA analysis was performed as described in the Supporting Information section IV.3.1. The sPCA values of each individual were converted to a color by translating each of the three principal scores into a color channel (red, green, and blue for 1^st^, 2^nd^ and 3^rd^ principal components respectively). To legibly display all samples analyzed, each individual sequence was placed as closely as possible to its original geographical location while avoiding overlaps. Different symbols were used to represent modern, historical and archeological samples. Additional samples for which only partial sequences were obtained but which contained enough information to allow unambiguous assignment to a specific clade are represented using a dotted outer line, assigning them the average color of the clade members. The specimens from the Hai-Bar Yotveta reserve in Israel that descended from hemiones captured in Turkmenistan and Iran are represented in the magnified box on the lower left side of the map connected by an arrow to their original location. Fig. S9 recapitulates the geographical location of samples

The sPCA scores, RGB and GPS values for each individual shown in the phylogenetic tree are listed in table S4. The sPCA scores correspond to the first three axes, as well as the last (36^th^) one. The RGB values are derived from an analysis that took into account only the first three axes. The graphical representation of the various results of the sPCA analysis are represented in Fig. S2-S6.

#### Median Joining Network

The median joining network of mitochondrial HVR (Fig. 2A) was constructed using the Network 4.6.0.0 program with a maximum parsimony post processing step (http://www.fluxus-engineering.com) [35].

**Figure 2:**
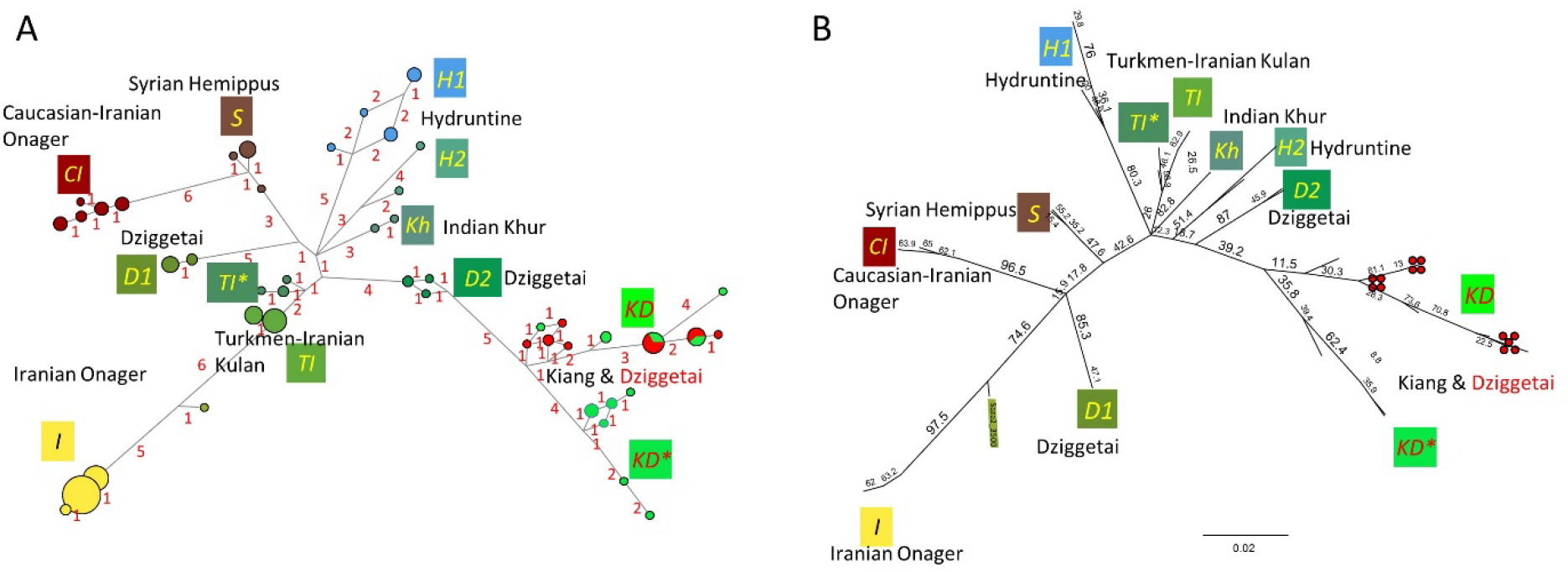
Median Joining Network analysis (A) and maximum likelihood analysis based on PHYML (B). The names of the deduced clades are indicated in italics. The colors of the box with the clade name follow the same convention as in the sPCA analysis displayed in Fig. 1.

#### Maximum Likelihood analysis

Maximum likelihood (ML) analyses (Fig. 2B) were computed using PHYML 3.0 [36,37]. Since *E. hemionus* mitochondrial sequences harbor a specific 28-bp-deletion that distinguishes them from all other equid sequences, and since the phylogenetic information of a deletion is not properly accounted for by substitution models, we performed ML analyses of the hypervariable regions using only sequences with this deletion. Based on model comparison criteria performed with jModelTest [38], we considered a TN93 model for the nucleotide substitution model [39] and a gamma-distributed rate of variation among sites (+G) with four rate categories (i.e., TN93+G model). We used 1000 bootstraps to estimate node robustness. We also used RaXML 8.2.3 [40] with the bootstrap convergence criterion autoMRE, which performed about a 1000 bootstraps, to determine the ML bootstrap support of the nodes of the maximum clade credibility tree of the Bayesian analysis presented in Figure S7.

#### BEAST analysis

Phylogenetic analyses were conducted under the Bayesian framework implemented in the program BEAST v. 1.7.5, which allows estimation of mutation and population history parameters simultaneously from temporally spaced sequence data [41]. We used the mitochondrial reference sequence of the domesticated donkey as an outgroup, and enforced monophyly to all sequences harboring the specific 28-bp-deletion. We used a strict molecular clock with a uniform prior over the (10^-12^,10^-6^) interval for the substitution rate measured in substitutions per site and per year. Based on model comparisons criteria performed with jModelTest [38], we further considered a TN93 model for the nucleotide substitution model [39] and a gamma-distributed rate of variation among sites (+G) with four rate categories (i.e., TN93+G model). Default priors were used for the seven parameters (alpha, kappa1, kappa2 and the four nucleotide frequencies) of the TN93+G nucleotide substitution model. Finally, a standard coalescent model was considered for the tree prior with a constant population size [42,43] and a log-normal prior distribution for the tree prior height with a mean of 13.5, a standard deviation of 0.843 and an offset of 1,500,000 years. This prior distribution is centered on 2,230,000 years, which integrate the various assumed and estimated divergence times between the ancestors of African and Asiatic wild asses [19,44,45], but within a relatively large range of possible values as the 95% credibility interval covers 1,640,000 to 5,300,000 years. The simplifying assumption of a constant population size overall was made to avoid overfitting of the data with too many parameters because populations must have expanded and contracted locally in a complex manner given the very wide spatial and temporal ranges considered in this study.

To estimate the posterior distribution of each parameter of interest, we used the Markov Chain Monte Carlo algorithm implemented in the BEAST software. We ran ten independent chains with initial values sampled as described above and an input UPGMA tree constructed using a Juke-Cantor distance matrix. Each of these chains was run for 10,000,000 iterations and for each parameter of interest, 4,500 samples (one every 2,000 generated ones) were drawn after discarding a 10% burn-in period. The BEAST output was analyzed with the software TRACER v. 1.5.0 [46]. Visual inspection of the traces and the estimated posterior distributions suggested that each MCMC had converged on its stationary distribution. In particular, effective sample size (ESS) values varied from around 300 for the kappa parameters of the nucleotide substitution model to around 3,000 for the tree height (most being over 600). Using Logcombiner, we further combined all the results from the 10 independent chains leading to combined ESSs ranging from 3,500 for the kappa parameters to 34,000 for the tree height. The maximum clade credibility tree with the median height of the nodes was finally calculated using TreeAnnotator V. 1.7.5 and visualized using FigTree v.1.4.0 http://tree.bio.ed.ac.uk/software/figtree/webcite.

#### Summary Statistics

The various summary statistics were computed using DNASP v5.1 [47] and Arlequin v3.5.1.3 [48] and are presented in Tables 1-3.

**Table 1:** Population pairwise Fst

Population pairwise FSTs: Distance method pairwise difference
|  | Gobi | Tibet | Iran (ancient) | Iran-Turkmenistan (modern) | Caucasus | Anatolia-Balkans | Syria |
| --- | --- | --- | --- | --- | --- | --- | --- |
| Gobi | 0 |  |  |  |  |  |  |
| Tibet | 0.14212 | 0 |  |  |  |  |  |
| Iran (ancient) | 0.34288 | 0.52891 | 0 |  |  |  |  |
| Iran-Turkmenistan (modern) | 0.42786 | 0.54144 | 0.29414 | 0 |  |  |  |
| Caucasus | 0.54529 | 0.73930 | 0.10069* | 0.57956 | 0 |  |  |
| Anatolia-Balkans (with Artenac) | 0.42281 | 0.60104 | 0.50186 | 0.55531 | 0.79964 | 0 |  |
| Syria | 0.45511 | 0.68097 | 0.33034 | 0.55005 | 0.82065 | 0.77230 | 0 |

FST P values (Number of permutations: 110)
|  | Gobi | Tibet | Iran (ancient) | Iran-Turkmenistan (modern) | Caucasus | Anatolia-Balkans | Syria |
| --- | --- | --- | --- | --- | --- | --- | --- |
| Gobi | * |  |  |  |  |  |  |
| Tibet | 0.00901 ± 0.0091 | * |  |  |  |  |  |
| Iran (ancient) | 0.00 ± 0.00 | 0.00 ± 0.00 | * |  |  |  |  |
| Iran-Turkmenistan (modern) | 0.00 ± 0.00 | 0.00 ± 0.00 | 0.00 ± 0.00 | * |  |  |  |
| Caucasus | 0.00 ± 0.00 | 0.00 ± 0.00 | 0.08108 ± 0.0212 | 0.00 ± 0.00 | * |  |  |
| Anatolia-Balkans (with Artenac) | 0.00 ± 0.00 | 0.00 ± 0.00 | 0.00 ± 0.00 | 0.00 ± 0.00 | 0.00 ± 0.00 | * |  |
| Syria | 0.00 ± 0.00 | 0.00 ± 0.00 | 0.00 ± 0.00 | 0.00 ± 0.00 | 0.00901 ± 0.0091 | 0.00 ± 0.00 | * |

For Iran, sequences from ancient and modern specimens are treated separately, as indicated.
Within the KD clade, using the maximum sequence length available for the HVR for the corresponding specimens (296 nt), the Fst value between kiangs and dzigettais is 0.2018 (Pval=0.00901 ± 0.0091).

**Table 2:** Pairwise Nucleotide divergence with Jukes and Cantor, K(JC) calculated with DNAsp

|  | Gobi | Tibet | Iran (ancient) | Iran-Turkmenistan (modern) | Caucasus | Anatolia-Balkans | Syria | India |
| --- | --- | --- | --- | --- | --- | --- | --- | --- |
| Gobi | * |  |  |  |  |  |  |  |
| Tibet | 0.03838 | * |  |  |  |  |  |  |
| Iran (ancient) | 0.06510 | 0.07351 | * |  |  |  |  |  |
| Iran-Turkmenistan (modern) | 0.05980 | 0.06110 | 0.05007 | * |  |  |  |  |
| Caucasus | 0.07209 | 0.08470 | 0.03220 | 0.05909 | * |  |  |  |
| Anatolia-Balkans (with Artenac) | 0.05641 | 0.05804 | 0.06664 | 0.05944 | 0.07527 | * |  |  |
| Syria | 0.05459 | 0.06275 | 0.04255 | 0.05217 | 0.03669 | 0.05851 | * |  |
| India | 0.05259 | 0.05806 | 0.05511 | 0.05469 | 0.05856 | 0.03756 | 0.04114 | * |

Within the KD clade, using the maximum sequence length available for the HVR for the corresponding specimens (296 nt), the pairwise nucleotide divergence between kiangs and dziggetais is 0.02294.

**Table 3:** Intrapopulation Diversity calculated using Arlequin and DNAsp

| Population | Nind | Nhap | Haplotype diversity | Nucleotide diversity, Pi | Theta(k) | Theta(S) | Theta(Pi) | Theta (per site) |
| --- | --- | --- | --- | --- | --- | --- | --- | --- |
| Gobi | 23 | 12 | 0.93 ± 0.03 | 0.041 ± 0.021 | 9.41 | 6.23 ± 2.36 | 9.40 ± 5.00 | 0.028 |
| Tibet | 17 | 11 | 0.95 ± 0.03 | 0.026 ± 0.014 | 12.40 | 4.44 ± 1.88 | 5.93 ± 3.33 | 0.019 |
| Iran (ancient) | 12 | 9 | 0.94 ± 0.06 | 0.043 ± 0.024 | 14.69 | 9.27 ± 3.89 | 9.89 ± 5.49 | 0.040 |
| Iran-Turkmenistan (modern) | 45 | 7 | 0.76 ± 0.05 | 0.028 ± 0.015 | 2.08 | 5.49 ± 1.87 | 6.51 ± 3.48 | 0.024 |
| Caucasus | 5 | 2 | 0.60 ± 0.18 | 0.008 ± 0.006 | 0.69 | 1.44 ± 1.02 | 1.80 ± 1.44 | 0.006 |
| Anatolia-Balkans with Artenac | 9 | 5 | 0.83 ± 0.10 | 0.018 ± 0.011 | 3.83 | 4.42 ± 2.16 | 4.11 ± 2.56 | 0.019 |
| Anatolia-Balkans without Artenac | 8 | 4 | 0.79 ± 0.11 | 0.012 ± 0.008 | 2.50 | 2.31 ± 1.31 | 2.79 ± 1.87 | 0.010 |
| Syria | 6 | 3 | 0.60 ± 0.21 | 0.005 ± 0.004 | 1.70 | 1.31 ± 0.91 | 1.20 ± 1.02 | 0.006 |

| Population | Nind | Nhap | Haplotype diversity | Nucleotide diversity, Pi | Theta(k) | Theta(S) | Theta(Pi) | Theta (per site) |
| --- | --- | --- | --- | --- | --- | --- | --- | --- |
| Dziggetai | 13 | 7 | 0.87 ± 0.07 | 0.015 ± 0.009 | 5.43 | 3.22 ± 1.53 | 4.51 ± 2.67 | 0.012 |
| Kiang | 17 | 12 | 0.96 ± 0.03 | 0.023 ± 0.013 | 16.80 | 5.32 ± 2.20 | 6.90 ± 3.82 | 0.018 |

## Results

### Population Structure of the Wild Ass in Asia and Europe

The data generated were subject to phylogenetic analyses, as well as to a spatial principal component analysis (sPCA; [33]), a multivariate method that investigates the potential relationship between geographic distances and genetic variation using a principal component analysis integrating genetic data and georeferenced positions of the samples (Fig. 1; see also Material & Methods and Table S4). For representing the scores of sPCA of each individual, the three principal scores were converted into a channel of color (red, green, and blue for 1^st^, 2^nd^ and 3^rd^ principal components, respectively), thus defining the color of each sample represented on a map (Fig. 1). The code color obtained through sPCA was also used to represent the samples in the other genetic and phylogenetic analyses: representation of the relationships between sequences with the median joining network (MJN, Fig. 2A), as well as maximum likelihood (ML, Fig. 2B, 4) and Bayesian (Fig. 3; Fig. S7) phylogenies.

**Figure 3:**
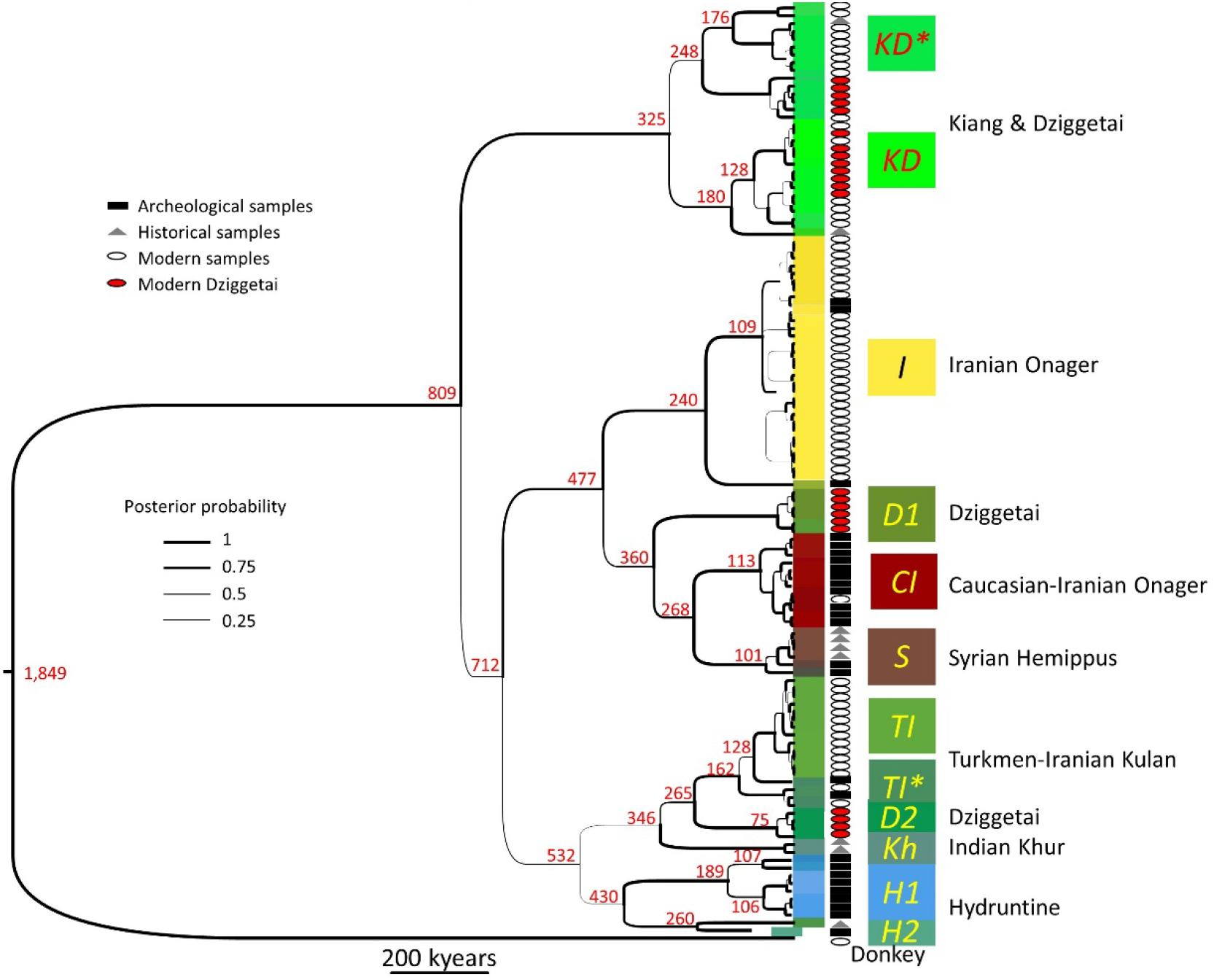
Phylogenetic tree of the mitochondrial control region of the Eurasiatic wild ass constructed through BEAST analysis. The corresponding *E. asinus* DNA region was used as an outgroup. The estimated median height of the nodes is indicated in red, in kyears, and the thickness of the lines is proportional to the posterior clade probability (the scale is represented). The mean substitution rate averaged across the whole tree is 8.5 E-8 substitutions per site per million years (95% HPD interval: 2.1-18.8 E-8). The colors of the box surrounding each individual sequence follow the same convention as in the sPCA analysis displayed in Fig. 1. The names of the deduced clades are indicated in italics. The symbols following each sequence name indicated the origin of the sample (Square: Archeological; Triangle: Historical; Circle: Modern), and the red circles indicate the modern dziggetais (see text). For an enlarged representation of the tree containing the names of the sequences, the 95% HPD of the node height values, the posterior probabilities of the nodes and their bootstrap values by ML analysis, see Fig. S7.

The sPCA reveals a clear phylogeographic structure of the data. The MJN and phylogenetic analyses show that the sequences recovered belong to nine clades (Fig. 2B and 3). Each of the nine clades is essentially dominant in a distinct geographical territory, i.e., Anatolia-Balkans, Syria, the Caucasus, the Tibetan Plateau, modern Iran-Turkmenistan and Northwest India, contrasted by ancient Iran and the Gobi with three different clades each. In the following, we will consider only the most robust phylogenetic relationships between clades that were consistently observed irrespective of the analysis method used.

We used summary statistic approaches to analyze the genetic diversity within and between each territory (Tables 1-3). The analysis of the genetic distance between the populations as expressed through the fixation index Fst is reported in Table 1. While there is moderate genetic differentiation between the modern populations from the Gobi and the Tibetan plateau (Fst=0.104), and between the ancient Caucasian and Iranian populations (Fst=0.1), the other populations including the modern wild asses from Iran and Turkmenistan, as well as the ancient populations from Syria, Anatolia and the Balkans are highly differentiated with Fst values between 0.33 and 0.82, a differentiation with high statistical support (Table 1).

Several measures of molecular genetic variability within populations were used, namely nucleotide diversity Pi (π) and the population parameter Theta (Θ) as estimated using several methods (Table 3). The highest genetic diversity was detected in the ancient Iranian and extant Gobi populations followed by the extant population of the Tibetan plateau (Table 3). The other populations are less diverse, the least diverse being the ancient, extinct populations from Anatolia and the Balkans, the Caucasus and Syria (Table 3).

In the following we describe the various clades grouped by larger geographical regions and correlate them with present-day subspecies.

***I, CI* and *TI* clades: onagers and kulans.** The ancient and modern Iranian specimens were found to belong to three clades (Fig. 2 and 3) named *I, CI* and *TI* that are distributed over a large portion of the phylogenetic trees (Fig. 2-3).

Among the three Iranian clades, the *CI* clade was once successful in the Caucasus before experiencing a severe decline during the last 4,000 years, with a single identified present-day representative of this clade corresponding to a wild ass with Iranian provenance [45].

A few ancient and most present**-**day Iranian onagers (*E. h. onager*) belong to the *I* clade whereas the present-day Turkmen kulans (E. *h. kulan*) belong to the *TI* clade (Fig.2 and 3). A small number of both ancient and modern Iranian onagers also belong to the *TI* clade, albeit to a sub-branch (*TI\**) that diverged from Turkmen kulans at an early stage of the radiation of the *TI* clade. The wild ass colony that was established in Israel in the 1970’s from 6 Iranian and 5 Turkmen wild asses belong to both the *I* and *TI* clades.

***Kh* clade: khurs.** The wild asses of Northern India, at present found in the reserve of the Rann of Kachchh (Kutch) and its surroundings, are called khurs (E. *h. khur).* We analyzed 19^th^ century museum specimens of the khur originating from Northwest India. Their sequences form the *Kh* clade. HVR sequences (240 bp) from the present-day khurs of Rann of Kachchh [49] share with the historic khur samples the characteristic SNPs of the *Kh* clade. The *Kh* clade is related to the Turkmen *TI* clade (Fig. 2 and 3).

***KD, D1* and *D2* clades: kiangs and dziggetais.** Mongolia and Tibet host two populations of wild asses: dziggetais (also called “kulans”; *E. h. hemionus*, a.k.a. *E. h. luteus*), and kiangs (*E. kiang).* The intrapopulation nucleotide diversity of the Mongolian dziggetais is the highest of all present-day wild ass populations, similar only to that found in the ancient Iranian population (Table 3). This high diversity suggests that these populations thrived in Mongolia and have not experienced bottlenecks as severe as those experienced by the other modern populations. All dziggetais studied belong to either the *KD, D1 or D2* clades (dziggetais are indicated by a red dot in Fig. 2-4), whereas those of the 17 analyzed kiangs belong exclusively to the *KD* clade. The branches of the *KD, D1 and D2* clades emerge at different locations in the phylogenetic trees (Fig. 2-3). These results suggest that there were three temporally distinct colonization events, most likely through Kazakhstan and the Dzungarian basin, the only route left accessible by the Himalayas. Within the KD clade, a KD* sub-clade can be distinguished that encompasses only kiangs, including the wild kiangs from the Southern part of Tibet (Fig. 4). The dziggetais from the KD clade have sequences that are closely related to the other kiangs analyzed, in particular to 80% of the kiangs originating from zoos (Fig. 4). The diversity of these sequences and the intricacy of their relatedness and identity (Fig. 4) argue that the presence of the *KD* clade in dziggetais is due to multiple events of mitogenome introgression that occurred over time from the kiangs to the dziggetais, rather than a single mixing event. Indeed, a large part of the diversity of the *KD* clade is found among both the kiangs and the dziggetais, including recently evolved haplotypes (compare the distribution of the dziggetais indicated by a red dot with that of the kiangs in Fig. 2 and 3). These admixture events must have been asymmetric, because 10 dziggetais belong to the *D(1+2*) and 13 to the *KD* clade, but none of the 17 kiangs belonged to either the *D1 or D2* clade. A Fisher exact test indicates that there is only a probability of 0.2% that such an unequal distribution would be observed in the absence of asymmetric gene flow. Thus, while it seems evident that some kiangs (at least females) have migrated from the Tibetan plateau to the Mongolian plain and interbred with dziggetais, we see no evidence that the reverse migration has occurred. Since mitochondrial sequences from geolocalized wild kiangs are only available from the Southern Tibetan population, from a region that is far removed from the area of distribution of the dziggetais (see Fig. 4), we cannot identify which kiang population(s) interbred with the dziggetais. We can only speculate that it involved (a) population(s) from northern Tibet with access to the Gobi. These recent and multiple admixtures question the validity of the classification of the kiang as a separate species (e.g., [50]).

**Figure 4:**
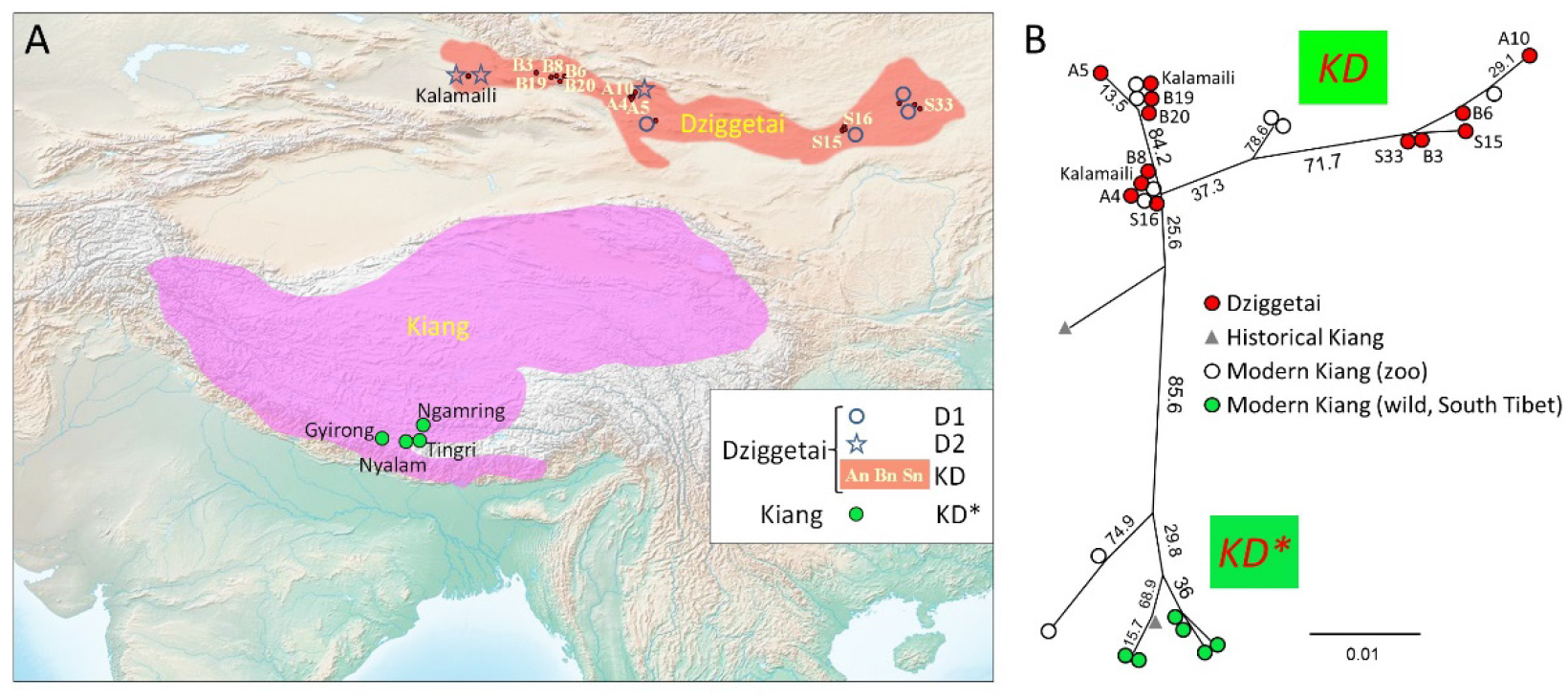
Geographic distribution of the analyzed kiangs and dziggetais and maximum likelihood phylogeny of the *KD/KD\** clade. **(A)** The map indicates the current areas of distribution of kiangs (pink) and dziggetais (orange) as determined by IUCN [6,8], as well as the location of the collected wild dziggetais (samples An, Bn, Sn; [73]; and the Kalamaili natural reserve; [81]) and kiangs (Nyalam, Gyirong, Ngamring and Tingri; [81]). Differences in the East-West distribution of the Dziggetai *D1* and *D2* clades are observed, although it is unclear how these correlate with the East-West distributions observed for microsatellite markers in [73]. **(B)** The ML phylogeny of the kiang and dziggetai sequences (295 bp) of the *KD* and *KD\** clades was performed using PHYML [36]. The bootstrap values of the nodes are indicated (1000 bootstraps). The modern and historical kiangs that were not geolocalized originate from zoos and museums, respectively.

**Clade *S*: hemippi.** The smallest of the Eurasiatic wild asses, the Syrian wild ass, or hemippus (*E. h. hemippus*), which stood only one meter at the withers, is now extinct. The last recorded animal was captured in the desert of Alep and died in 1929 in Vienna (Fig. S13) [51]. Expectedly, the mitochondrial lineages from four museum samples of the 19^th^ and 20^th^ century cluster together but, strikingly, these same lineages are also found in ancient specimens from Tall Munbāqa (Syria), located on the Middle Syrian Euphrates and dating to ca. 1,500-1,200 BCE. Together, they form the *S* clade that is related to the *CI* and *D1* clades. The paternal lineage of the hemippus based on the analysis of the Y chromosomes of both ancient and museum samples is also distinct from that found in other wild asses (Bennett et al., in preparation), indicating a continued reproductive isolation of this group.

Our observation of the genetic continuity over 5,000 years of the Syrian wild ass population led us to revisit the observation that Syrian wild asses from the site of Shams ed-Din Tannira (6^th^ millennium BCE) were larger than the modern hemippi [52]. We thus compared osteometric data from the Shams ed-Din Tannira specimens with those of this study: the 19^th^-20^th^ century hemippi, and Bronze Age Syrian wild asses from Tell Munbāqa (2^nd^ millennium BCE). For this comparison osteometric data (provided by L. Gourichon and D. Helmer) from 10^th^-9^th^ millennia BCE Tell Mureybet, located in the middle Syrian Euphrates valley, was also used, and, as an outgroup, individuals from Göbekli Tepe (10^th^-9^th^ millennia BCE), about 150 km further north in Southeast Anatolia, where genetically determined individuals belonged to clade *CI*. The comparison of various measurements from postcranial skeletal parts reveals that the prehistoric Syrian individuals range within the upper part of the size variation of their modern descendants or even surpass these in size (Supporting Information, section III.1.4.; Table S11 and Fig. S11A-B). This analysis showed a difference in average bone size between the prehistoric Syrian and Anatolian sub-populations: the wild asses hunted near Göbekli Tepe were clearly heavier than those living further south (Tell Mureybet and Munbāqa). The difference from these two areas both in bone size and of the two corresponding mitochondrial clades point to two distinct morphotypes. Thus, even though the prehistoric Syrian wild asses were already smaller than their neighboring Anatolian relatives, they were still of a more robust build compared to their modern descendants. The new genetic evidence presented here thus supports the hypothesis that the animal became smaller in the relatively recent stages of its evolution preceding its extinction [52].

***H1* and *H2* clades: hydruntines.** After observing inconsistencies between our preliminary genetic data and the initial taxonomic assignment of certain remains, we subjected all available bone and tooth samples from which we obtained DNA sequences to a “cross-determination” performed as a blind test by several of the osteologists participating in this research. In a number of cases, substantial disagreement among them was observed (Table S14). As emphasized previously ([5] and citations therein), this demonstrates the difficulties of assigning equid remains to (sub)species level based on osteomorphology and/or osteometry. For this reason, paleontologists tend to include non-morphological criteria in their taxonomic assessments, such as the time period and the geographical area from which the specimen in question originated, a strategy which can lead to circular reasoning (see discussion in [5]. Accurate taxonomic identification of hydruntine (*E. hydruntinus*), known only from fossil remains for which osteological and odontological diversity has not been well characterized, proved to be particularly problematic. In our study, 40% of the bones and teeth assigned on osteomorphological grounds to hydruntine by a subset of the osteologists involved yielded sequences clustering with *E. caballus.* Some of the caballine sequences obtained from these ancient specimens are either at present extinct or have not yet been found in modern horses (data not shown). Thus, for this taxon we adopted a ‘specificity index’ based on the degree of agreement among the specialists of equine osteomorphology that allowed us to consider archeological specimens most confidently assigned as hydruntine (See discussion in the Supporting Information section III.3, Fig S11 and Table S14). This procedure enabled us to characterize the distinct genetic structure of these ancient populations that has eluded previous attempts.

This approach revealed an extinct clade, *H1*, comprising sequences from archaeological specimens dated from 5,000 to 8,000 years BP, from bones and teeth that achieved the highest specificity index in our classification of morphological determination of hydruntine (Fig. S12; Table S14). This group comprises all remains that were identified as hydruntine with a specificity index > 0.5 including the three remains all osteologists unanimously agreed upon. Nearly all specimens found in Anatolia and the Balkans belong to the *H1* clade (Fig. 2-3). Despite their geographical proximity, the *H1* clade is genetically distant from the *S, CI* and *I* clades and appears more closely related to the *TI, Kh* and *D2* clades suggesting a different phylogenetic history than that of its geographic neighbors. Additionally, a metapodial from the Early Upper Pleistocene cave of Artenac, France, with a stratigraphic age of roughly 100,000 years was determined as clearly belonging to hydruntine with a specificity index of 1 (see Table S14 and Fig. S12). This sample yielded a mitochondrial sequence that was assigned to a distinct H1-related clade, *H2.* Surprisingly, this clade was also found in an early 20^th^ century museum specimen originating from Iran. This allowed us to identify two mitotypes: *H1* and *H2*, most likely representing the extinct hydruntine. All presumed hydruntines that belonged genetically to another wild ass clade had an average specificity index at least twice as low (Fig. S12). The phylogenetic proximity of the *H1* and *H2* clades with the *Kh, TI* and *D2* clades (Fig. 2B and 3) indicates the hydruntine to be a subspecies of *E. hemionus*, to the same degree as kiangs, dziggetais, hemippi, kulans, khurs, extinct and extant onagers, altogether forming a group that can be referred to as the Eurasiatic wild ass. We propose therefore to classify it as *Equus hemionus hydruntinus.*

## Discussion

### Origins

Equids evolved in North America before dispersing into Asia around 2.6 million years ago (Mya) [53]. Genomic data indicate that caballoid and stenonid equids diverged from each other ca. 4.5 Mya, and that gene flow between them ceased around the beginning of the Asian dispersal [16]. Among stenonids, the separation of asses and zebras has been estimated at ca. 2 Mya while that of African and Eurasiatic asses has been estimated at ca. 1.7 Mya ([16], see Fig. S8). Our data, based on the mitochondrial HVR, are consistent with these estimates (Fig. S8). Since the mitochondrial genomes of all Eurasiatic wild asses are characterized by a 28-bp deletion that has a very low probability of being homoplasic, they most likely constitute a monophyletic group. Where did this group emerge? All present-day stenonids are African, except for the Eurasiatic wild asses, and the earliest datable fossil evidence of *Equus* in Africa occurs ca. 2.33 Mya [54]. Thus, it could be hypothesized that the last common ancestor of all present-day stenonids was African. In this scenario, the Eurasiatic wild ass could have emerged ca 1.7 Mya from an ancestral ass-like population living in northern Africa, the Arabian peninsula and the Levant. Since all Eurasiatic wild ass mitogenomes evolved from an ancestral stenonid mitogenome after a single 28-bp-deletion event has occurred, the ancestral population of the Eurasiatic wild ass must have gone through a severe bottleneck before or during its migration into Eurasia. Alternatively, the ancestral stenonids population could have evolved in the southern plains of Asia, from where at least two independent migration waves into Africa gave rise first to the zebras around 2 Mya, and second to the African asses ca. 1.7 Mya. In this scenario, a severe bottleneck must have affected the population that remained in Asia leaving only the descendents of the deletion-harboring lineage that emerged between about 700 and 800 kiloyears ago (kya) ([16,45] and Fig. 3). The mitogenome sequence of a ca. 45,000-year-old Siberian *Sussemionus (E. ovodovi*) is in favor of this latter hypothesis since it reveals an extinct Asian lineage that has diverged from other stenonids around the time of separation of the zebras and the asses ([45]; Fig. S8). Genomic data do not yet enable us to decide which of the two hypotheses is more likely, and fossil evidence in certain regions is lacking, due in part to taphonomic reasons, but also to the rarity of detailed morphological description, consistent analysis and rigorous comparison [54]. Following divergence from other stenonids, the ancestral population of hemiones would have dispersed on the Eurasiatic continent where the populations would have further evolved and phylogeographic stratification taken place.

The phylogenetic relationships between the various mitogenome clades do not reveal a simple relationship with geographical distance but rather suggest a complex phylogeographic history with back-and-forth migrations. Our data suggest that Eurasiatic wild asses harboring the *KD/KD\** clade mitogenome may have migrated during the Middle Pleistocene into the Gobi and Tibet where they evolved independently. Eurasiatic wild asses with either of the *I, CI, S, D1* clade mitogenomes may have evolved in Southwest Asia where most of them were found in the Holocene, from where some of them (*D1* clade) migrated to the Gobi, presumably during the end of the Middle Pleistocene. Central Asia, where clade *TI* is still found, may also have allowed evolution of the Eurasiatic wild asses that are related to the *TI* clade and that have spread into Europe (*H1* and *H2* clades), India (*Kh* clade) and the Gobi (*D2* clade). Since the *H1* and *H2* clades are not closely related to the clades established in Southwest Asia (I, *CI*, S), we hypothesize that they have colonized Europe during the Pleistocene through a Northern route involving the Pontic-Caspian Steppes and that they arrived later in Anatolia coming from Europe at times when the Bosphorus was a land bridge. Such a scenario would explain the strong differentiation of the *H1* clade in Anatolia with respect to the geographically neighboring populations of the Syrian hemippi *S* and the Iranian onagers *I* and *CI* as well as their closer relatedness to the Turkmen kulans *TI*, the Indian khurs *Kh* and the Mongolian dziggetais *D2* (Fig. 1-3).

### Taxonomy and Conservation Biology

Conservation programs aim to preserve the evolutionary potential of a species using the classification of populations by their evolutionary significance based on ecological, morphological, geographic and genetic criteria [15,55]. The characterization of clades presented in this study thus provides a helpful guide for taxonomy and conservation biology. Our dataset reveals events of past and recent mitochondrial introgression between populations that are now considered separate species, such as kiang (*E. kiang*), or subspecies, such as onager (*E. h. onager*) and kulan (*E. h. kulan*) [50]. Poor genetic differentiation between kiangs and dziggetais (*E. h. hemionus*) has also been observed in a microsatellite study of equid diversity involving a smaller sample size (6 kiangs and 3 dziggetais) [56], indicating that our observation is not a peculiarity of the mitogenome transmission. We believe it is more appropriate to consider the kiang as a distinct population or perhaps even a metapopulation [57] of *E. hemionus*, with specific adaptations to the high-altitude climate and vegetation of the Tibetan plateau.

The designation of onagers and kulans as separate evolutionary significant units has been questioned [58]. Among the three clades that had representatives in Iran during the last 8,000 years, the *I* clade remained centered in Iran and is the prevalent clade in present-day onagers; the *CI* clade shows a cline towards the Caucasus, where the corresponding population is now extinct, and the *TI* clade shows a cline towards Turkmenistan. All three clades coexisted in the past at a single location near present-day Tehran (Sagzabad, 3,500 ya), and members of these clades are still interbreeding, showing that these clades do not define true diverging allopatric lineages. Currently, Iranian and Turkmen wild asses kept in the Hai-Bar Yotveta reserve in Israel are reported to interbreed and hybrids thrive without showing signs of outbreeding depression [59] [60]. Given the fact that these populations are shrinking rapidly, it is worth considering that in a not so distant past, when they occupied large interconnected areas, crosses between neighboring populations allowed gene flow events that have only recently been interrupted, enhancing the risk of inbreeding depression. Ensuring the survival of the Asiatic wild ass is a challenge that may justify managing the last remaining populations as components of a viable metapopulation [61].

### Palaeoecology of the Eurasiatic wild ass

The repeated glaciations that alternated with warmer phases throughout the Pleistocene had major impacts on the fauna, flora and the environment (e.g., [1,62]). These climatic oscillations were likely to also affect the distribution, speciation and population size of the wild asses. Indeed, in Western Europe, hydruntines were present only during the warmer and more humid interglacial periods of the Pleistocene (e.g., [12,63]). This Western European ecomorphotype was apparently adapted to milder climatic conditions and hilly landscapes (e.g., [12,63]). Indeed, the analyzed specimens from the caves of Artenac and Quéroy in Western France were dated to ca. 100,000 (the Eemian interglacial) and 12,000 ya, respectively, periods characterized by a milder climate corresponding to Marine Oxygen Isotope Stages 5 and 1 (e.g., [64-66]). The populations of *E. hydruntinus* adapted to the warmer and more humid climate in Europe during the interglacial stages were probably repeatedly separated from each other and/or went locally extinct during subsequent glacial periods [10], a process that was possibly accelerated through competition with cold-adapted horses [67]. For example, we identified an extant onager belonging to the same *H2* clade as the ca. 100,000-year-old individual from the Artenac cave in France might be a descendant of the populations that retracted to the Northern Middle East during the Lower Pleniglacial cold period roughly 70,000 ya (Fig. 3). Correspondingly, during the cold periods of the Pleistocene, the European wild ass likely withdrew to Southwest Asia, solely or in addition to the Southern European glacial refuges, a behavior we also observed for the European bison [1].

The genetic structure of the Asiatic population of the wild ass seems conditioned by geographical and climatic factors: the Asiatic steppe belt, the Iranian highlands and the Kara Kum desert in Turkmenistan, the mountainous Armenian highlands (Caucasus and Western Iran), the arid lowlands of Syria-Mesopotamia, the Anatolian highlands and the Balkans. Each of these ecogeographical units harbored a genetically distinct population, which therefore can be considered to be different ecomorphotypes. During the last glacial maximum (LGM), Anatolia’s forests and woodlands disappeared and were replaced by cold steppe vegetation [68], climatic conditions that could be compared to those of present-day Tibet where kiangs live today. Thus, it may have been a favorable habitat for the Asiatic wild ass. Colder periods in the Pleistocene and the beginning of the Holocene, when the sea level was low, would have allowed exchange between the European and the Anatolian populations until around 10,600 - 7,600 BP, the presumed date of the filling of the Black sea (e.g., [69]). The steppe vegetation remained dominant in Anatolia even after 10,000 BP, when humidity increased, slowly being replaced by woodlands, a change that reached its peak around 8,000 BP, followed by a last regression until 6,800 BP [68]. Unstable climatic conditions which began after 8,200 BP were marked by climatic oscillations and fluctuating hydromorphological conditions, leading to drought and heavy floods and to the increase of swamps in central Anatolia [70]. This, alongside anthropogenic pressure, could have been one of the reasons for the gradual disappearance of the Anatolian-Balkans population. The most recent specimens from the Anatolian plain harboring the *H1* mitotype date from 4,200-4,000 BP setting a lower limit for the date of disappearance of the hydruntine in Anatolia.

The differentiation of the Syrian population with respect to the Iranian and Anatolian populations could be explained by a long independent evolutionary history of these wild ass populations caused by the ecological barriers of the Zagros and the Taurus mountain ranges, which separate the Syrian plain from the plateaus in Anatolia and Iran. South of the Taurus Mountains, landscape and vegetation near the headwaters of the Upper Balikh received more precipitation than the Middle Syrian Euphrates Valley near Tell Mureybet and likely offered more favorable living conditions to *E. hemionus* [71]. This may explain the larger size of the Early Neolithic *E. hemionus* from Southeast Anatolian Göbekli Tepe compared to the contemporaneous specimens from Middle Syrian Tell Mureybet and Late Bronze Age Tell Munbāqa. Following the main vegetation zones [71], the larger subspecies was associated with the xerophilous deciduous forest steppe, while the smaller one was confined to the Mesopotamian steppes. Drought, desertification, overgrazing by livestock and increased anthropogenization of the landscape combined with hunting pressure may explain the decline and extinction of the hemippus. Despite genetic continuity through time, the Syrian wild ass population witnessed a significant and previously unreported reduction in body size preceding its extinction. This might be the result of impoverished living conditions and/or increased temperature in historical times pushing it into an ecological pessimum or the consequence of inbreeding depression due to a decrease in population size (see Supporting Information section III 1.4.1).

The Asiatic populations of the North-East (kiangs and dziggetais) and of the South-East (onagers, Turkmen kulans, khurs and hemippi) have been continually separated by the Central Asian massif, which was covered by glaciers during the cold periods [72]. Nevertheless, at least limited gene flow between populations of Eurasiatic wild asses could have taken place via the Irtych valley and the Dzungarian basin, which were accessible owing to a very dry climate [10]. The observation that the extant dziggetai *D1* and *D2* clades cluster with the ancient and present-day onagers and kulans argues in favor of the existence in the rather recent past of a population continuum with introgression from Iran and/or Central Asia to the Gobi. The other dziggetai *KD* clade indicates introgression from the kiangs on the Tibetan plateau to the dziggetai in the Gobi.

For the onagers and kulans, the remaining extant populations seem to represent endemic relict populations of previously much larger, connected populations living in the Caucasus, on the Iranian Plateau and in Turkmenistan.

In summary, in the past, the Eurasiatic wild ass was able to adapt to changing climatic conditions through population range shifts, which has become increasingly difficult due to habitat destruction and fragmentation, preventing the animals from reacting according to their natural behavior and local habitat conditions [73].

### *H1* and *H2*, the hydruntine mitotype

It was formerly assumed, based on geographic distribution through time, rate of speciation and capacity for sympatry deduced from morphological features of fossil remains, that stenonid and caballoid horses of the genus *Equus* differed ecologically, the stenonids being more specialized and therefore adapted to narrower niches [74]. In contrast, the ecological flexibility in the caballoids was considered to be the consequence of behavioral versatility rather than increased morphological variation [74]. The present results question this conclusion since fossils with presumed stenonid features were found during the course of this study to belong to the caballoid horses. This indicates that the morphological plasticity of past equids appears to be higher than previously assumed and that some criteria used to determine species within this genus are in fact pleiomorphic.

Despite these identification difficulties, we could obtain data from a sufficient number of consensually assigned hydruntine bones to establish that the *E. hemionus H1* and *H2* clades correspond to the paleontological species of *E. hydruntinus* (see section III.3 and table S14 of the supporting information). Perhaps due to incorrect taxonomic identification of samples, no hydruntine-specific mitochondrial signature had been found in previous paleogenetic studies, which considered hydruntines to be an onager-like wild ass [18,19]. The co-occurrence of an *E. hemionus* mitotype signature with a set of distinctive morphological features found in this study argues in favor of the hydruntine being a particular morphotype or ecomorphotype of *E. hemionus.* Since the separation of the *H1+2* clade from other mitotypes of *E. hemionus* is not as ancient as that separating mitotypes of interfertile populations, like the *KD* and *D* clades of the dziggetais, our data do not support the classification of the hydruntine as a distinct species. Instead, it is probably more appropriate to consider it a subspecies (E. *h. hydruntinus*), as has been proposed for other current Eurasiatic wild ass populations, even though it is not clear whether in biological terms this level of taxonomic differentiation corresponds to something more than naming a population. The identification of the hydruntine as a Eurasiatic wild ass finds additional support in contemporary cave art representations, such as in Lascaux cave (Fig. 5). The presence of *E. hemionus* in Europe when these works were created challenges popular assumptions that, due to a previously presumed absence of this species in Europe during the Upper Paleolithic, these images must therefore represent “deformed” horses [75]. Representations of the hydruntine in other French caves (Engraving in the cave “Les Trois Frères”, Grottes des Volpes, France; Engraving on a pendant in the cave of Putois, France) resemble present-day hemiones (Fig. 5B and C) but show even longer ears. Strikingly, long ears are also a distinct feature of the wild asses represented in hunt scenes on Late Neolithic vessels excavated from the Anatolian site of Kösk Höyük (Fig. 5D). In this site we found eight equid bones with H1 haplotype suggesting that these depictions are representations of the local hydruntine since there is no evidence for donkeys in Anatolia until the 4th millennium BC at the earliest (e.g., [76,77]). Altogether these representations suggest that the animals’ ears were characteristic enough to be depicted. Their similarity lends further support to our finding that the hydruntine populations from Anatolia and Europe were closely related.

**Figure 5:**
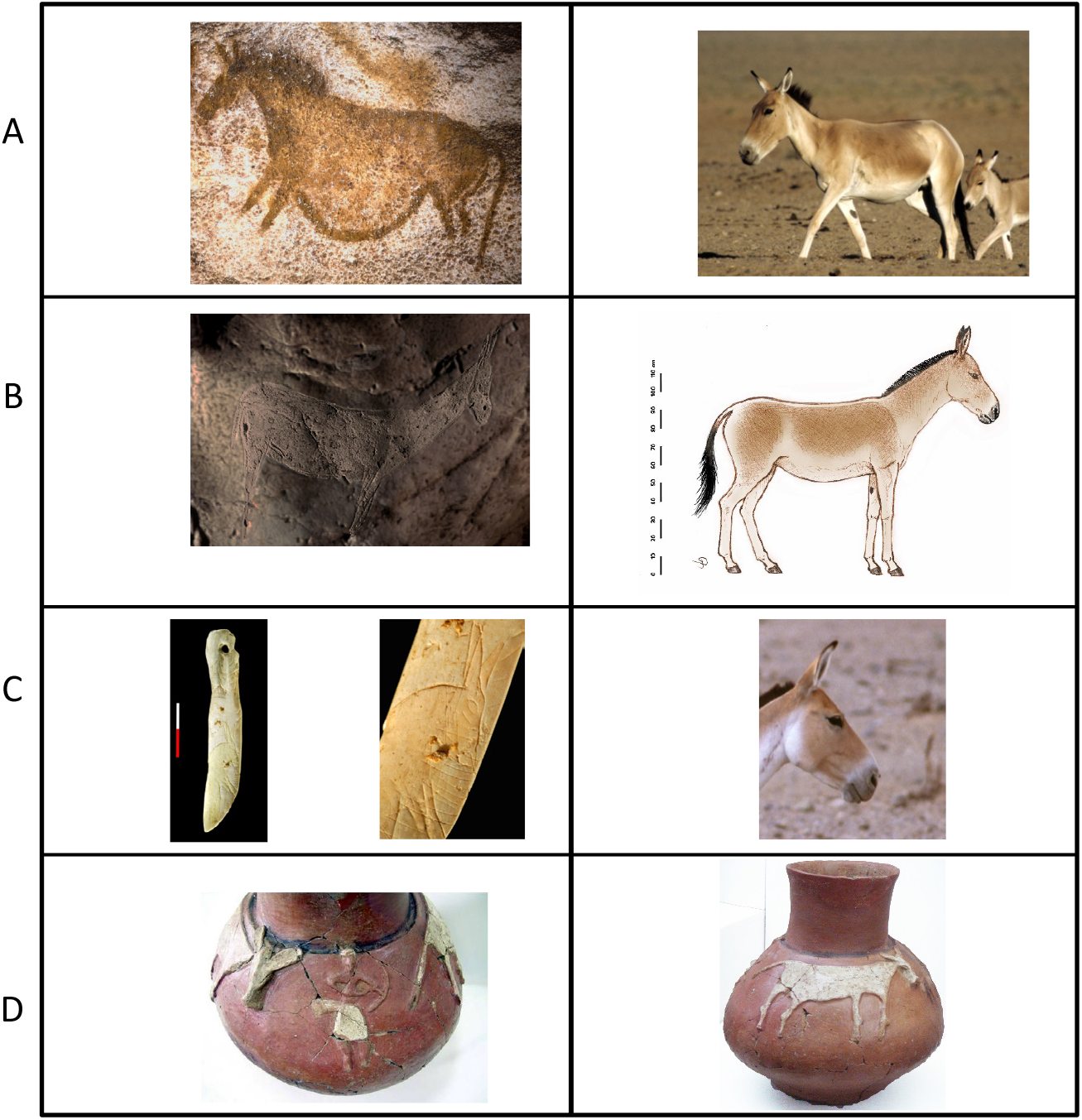
Ancient representations of the Eurasiatic wild asses. **A:** Comparison of the “Panel of the hemione” from the Lascaux cave, France (Copyright: N. Aujoulat CNP-MCC) with a Dziggetai from the Southeast Gobi in Mongolia (Photo & Copyright P. Kaczensky). **B:** Engraving of a presumed hydruntine in the cave “Les Trois Frères”, Grottes des Volpes, France (Photo & Copyright: Robert Bégouën), and graphical reconstruction of a hydruntine based on prehistoric engravings as well as the morphological and genetic results presented here (Drawing and copyright: Erich Pucher). **C**: Comparison of an engraving of a presumed hydruntine on a pendant in the cave of Putois, France (Photo & Copyright: © Musée National de Préhistoire MNP, Les Eyzies Dist.RMN - Photo: Ph. Jugie) with an onager from Touran, Iran (Photo & Copyright: Gertrud & Helmut Denzau). **D**: Depiction of a hunt scene of presumable hydruntines on a vessel from the Late Neolithic site of Köºk Höyük in Anatolia (Photo & Copyright: Aliye Öztan.

The surprising recovery of the *H2* subgroup known from a 100,000-year-old specimen from Western Europe, which may represent an older hydruntine population, from an early 20^th^ century museum specimen from Persia, suggests the possibility of ancient interbreeding events between hydruntines and other Eurasiatic wild asses. Two other specimens yielded incomplete sequences that were nevertheless sufficient to assign them to the *H1* subgroup, one from the Queroy cave in Western France, 12 – 13,000 years BP, and one from Scladina cave [18] in Belgium, estimated to be between 30,000 and 40,000 years old (D. Bonjean, pers. comm.). Later specimens from Romania and Turkey dated 8,000 to 4,000 years BP belong to the *H1* subgroup. Thus, all specimens belonging to the H1 subgroup, which shows a low level of diversity, are between 40,000 to 4,000 years old (see Table S1). The distance from the *H1* to the *H2* mitotype suggests that the hydruntine population could have gone through a bottleneck during the last two glacial periods and that Europe could have been recolonized from a refugial population, as proposed earlier on morphological grounds [12]. Notably, “hydruntine” specimens post-dating 4,000 years BP do not contain the genetic signature of hydruntines (Table S1). Our results are indicative of the disappearance of the hydruntine-type of Eurasiatic wild ass around the end of the Bronze Age, presumably following habitat fragmentation and human exploitation as proposed recently [13], but a pocket population might have remained on the central Anatolian plateau and on the Iberian peninsula until historical times [78].This characterization of a past population of Eurasiatic wild ass in Europe should be noted when considering low-intervention conservation management strategies of abandoned rural areas such as “rewilding”, where an abundant wild large herbivore population was concluded to be instrumental in maintaining biodiversity of vegetation structures under a temperate climate in the absence of human management [79]. Although the challenges of the reintroduction of species which have disappeared from Europe a few millennia ago are many [80], consideration of the Eurasiatic wild ass may be appropriate for such initiatives.

## Acknowledgments

We wish to express our respect to Véra Eisenmann and our gratitude for the taxonomic determination of ancient equid specimens. We are grateful to Cédric Beauval, Eric Boëda, Marie-Françoise Bonifay, Jean-Philip Brugal, Jean-Christophe Castel, Jean-Jacques Cleyet-Merle, Lydia Gamberi, Christophe Griggo, Joséphine Lésur, Stéphane Madelaine, Gabriella Magnano, Marylène Pathou-Mathis, Eric Pellé, Maryline Rillardon, Elodie Trunet, Frank Zachos, João Zilhão, for providing samples for our study, which did not yield genetic results, as well as to Thierry Petit for helping to sample onager feces; to the curators of the Musée National d’Histoire Naturelle, Paris, France, the Naturhistorisches Museum Wien, Vienna, Austria, and the Réserve Africaine de Sigean, Sigean, France for providing specimens and samples; to the studbook keeper Claus Pohle, Tierpark Berlin, Germany, and Thomas Lind, Animal Coordinator at the Kolmården Wildlife Park, Sweden for help with the studbook research; to Gertrud and Helmut Denzau for the photo from the Touran reserve in Iran, the Musée de Préhistoire, Les Eyzies, France, for the photos of the “pendeloque” of the cave of Putois, Robert Bégouën for the photo of the hydruntine carving from the cave of “Les Trois Frères”, and Elena Man-Estier from the Centre National de Préhistoire, Ministère de la Culture et de la Communication, Direction Générale des Patrimoines,for the photo of the “panel of the hemione” from the Lascaux cave.

## Data accessibility

*E. hemionus* HVR mitochondrial DNA sequences: Genbank: Accession numbers to be provided upon manuscript acceptance

## Author contributions

Conceptualization: EMG, TG

Data Curation: TG, EMG

Formal Analysis: TG, MGa, RK, JP

Funding Acquisition: EMG, TG, BSA, JP

Investigation: EAB, SC, JP, SG, MP, SBD, SJMD

Project Administration: TG, EMG

Resources: BSA, PK, RK, MM, AMM, HPU, AB, MGe, CYG, MRH, PEM, AÖ, EP, JFT, MU, CW

Supervision: EMG, TG

Validation: EMG, TG, JP, SJMD, HPU

Visualization: EMG, TG

Writing – Original Draft Preparation: EMG, TG, EAB

Writing – Review & Editing: EMG, TG, EAB, JP, BSA, SJMD, HPU, PK, MM, AMM, RK, MGe, CYG, JFT, PEM, MRH

## Supporting information

### List of tables

All tables are presented as separate tabs in a single Excel file

Table S1 Description of all samples analyzed

Table S2 Primers used to amplify mitochondrial sequences

Table S3 Description of published sequences used

Table S4 Sample location and results of sPCA analysis

Table S5 Population Pairwise FSTs

Table S6 Pairwise Nucleotide divergence with Jukes and Cantor, K(JC) calculated with DNAsp

Table S7 Intrapopulation Diversity calculated using Arlequin and DNAsp

Table S8 Nucleotide positions on which assignment to clades is based.

Table S9 Characteristics, measurements (mm) and zoological assignment of samples from Iranian archeological sites

Table S10 Comparison of measurements from the sample DAG2 from Daghestan-Velikent with those from other hemiones and a kiang

Table S11 Comparative measurements of skeletal parts of late and Bronze Age E. hemionus hemippus

Table S12 Measurements of the teeth of the hydruntines of Cheia

Table S13 Comparative measurements (mm) of the analyzed metacarpals of the caves of Artenac and Pair-Non-Pair

Table S14 Determination of the specificity index of the hydruntine bones

Table S15 Studbook register of the founder animals from the Hai-Bar-Yotveta Reserve

### List of figures

The figures are all grouped in a single supporting document alongside with the detailed description of the samples and of the archeological sites.

Figure S1: Global alignment of all sequences obtained and used for the various analyses

Figure S2: sPCA, distribution of the eigenvalues

Figure S3: sPCA, spatial and variance components of the eigenvalues (screeplot)

Figure S4: sPCA, distribution of the first three principal components of the sPCA for samples colored according to their origin

Figure S5: sPCA, histogram of the simulation to test the significance of the global structure

Figure S6: sPCA, histogram of the simulation to test the significance of the local structure

Figure S7: Phylogenetic tree of Hemione mitochondrial control region constructed through BEAST analysis.

Figure S8: Dating estimates of the equine phylogeny.

Figure S9: Geographical location and time periods of the samples that yielded genetic results

Figure S10: Geographical distribution of the different past and present wild ass populations in Europe and Asia and the principal present-day reserves where they still occur

Figure S11A: Osteometrical data from Syrian hemiones

Figure S11B: Logarithmic Size Index method

Figure S12: Histogram of specificity index and clade of bone samples determined as E. hydruntinus

Figure S13: E.h. hemippus

Figure S14: Dziggetais at a pothole in the Gobi. (Photo: P. Kaczensky)

FigureS15: Onager (right) and kiang (left) (Réserve Africaine de Sigean, France)

